# Failed stopping transiently suppresses the electromyogram in task-irrelevant muscles

**DOI:** 10.1101/2024.02.27.582386

**Authors:** Isaiah Mills, Mitchell Fisher, Corey Wadsley, Ian Greenhouse

**Affiliations:** Action Control Lab, Department of Human Physiology, University of Oregon, Eugene, Oregon, USA

## Abstract

Selectively stopping individual parts of planned or ongoing movements is an everyday motor skill. For example, while walking in public you may stop yourself from waving at a stranger who you mistook for a friend while continuing to walk. Despite its ubiquity, our ability to selectively stop actions is limited. Canceling one action can delay the execution of other simultaneous actions. This stopping-interference effect on continuing actions during selective stopping may be attributed to a global inhibitory mechanism with widespread effects on the motor system. Previous studies have characterized a transient global reduction in corticomotor excitability by combining brain stimulation with electromyography (EMG). Here, we examined whether global motor inhibition during selective stopping can be measured peripherally and with high temporal resolution using EMG alone. Eighteen participants performed a bimanual anticipatory response inhibition task with their index fingers while maintaining a tonic contraction of the task-irrelevant abductor digiti minimi (ADM) muscles. A time series analysis of the ADM EMG signal revealed transient inhibition during failed stopping compared to go response trials 150 ms to 203 ms following the stop signal. The pattern was observed in both hands during bimanual stop-all trials as well as selective stop-left and stop-right trials of either hand. These results indicate that tonic muscle activity is sensitive to the effects of global motor suppression even when stopping fails. Therefore, EMG can provide a physiological marker of global motor inhibition to probe the time course and extent of stopping processes.

**Key Points:**

- Successfully stopping an initiated response globally suppresses the motor system.
- Using electromyography of tonic muscle activity, we show inhibition spills over to task-irrelevant muscles during failed stopping.
- The electromyographic pattern of inhibition is transient, lasting from approximately 150 to 203 ms following a stop signal when stopping fails.
- The time course of the peripheral suppression of muscle activity may be leveraged to more precisely examine candidate neural mechanisms.
- This non-invasive measure of motor system inhibition may be useful for tracking inhibitory control deficits in clinical populations.

## Introduction

Response inhibition is the process of canceling planned or initiated actions to meet environmental demands. Often, these demands require us to stop only one part of our action plan while continuing the remaining parts. For example, while walking through a crowd, seeing somebody you know may cue you to raise your hand to greet them. If you realize the person is not who you thought they were, you may cancel the movements associated with a greeting while continuing to move your legs to walk. The ability to selectively stop a subset of actions is essential for flexible interactions with our environment. In the laboratory, behavioral stopping is often investigated using response inhibition tasks in which participants execute a response associated with a go signal and attempt to cancel this response upon the presentation of a stop signal (Logan and Cowan, 1984). Evidence from many studies using various iterations of stop-signal and anticipatory response inhibition (ARI) tasks indicates that stopping globally inhibits the motor system. In particular, corticomotor excitability (CME) is reduced in both task-relevant and -irrelevant muscles during stopping, as indexed by decreased motor evoked potential (MEP) amplitudes elicited with transcranial magnetic stimulation (TMS) (Badry et al., 2009; Majid et al., 2012). Global inhibition is generalizable, as stopping verbal responses (Cai and Aron, 2012) and inhibiting eye saccades (Wessel et al., 2013) also reduced CME of task-irrelevant hand muscles.

Global effects of response inhibition on the motor system are observable during selective stopping tasks which require the cancellation of part of a multi-effector response (Wadsley et al., 2022a). Selective stopping is typically investigated using multicomponent response inhibition paradigms during which a stop-signal is presented for only one effector while the remaining effectors should respond as intended. Selective stopping in these paradigms results in the delayed execution of the non-stopping effector, a phenomenon referred to as the stopping-interference effect (Coxon et al., 2007; Aron & Verbruggen, 2008; MacDonald et al, 2012; Wadsley et al. 2019). The stopping-interference effect is also attributable to a global inhibitory mechanism, as evident in suppressed CME of the non-stopping effector during the time of response inhibition (MacDonald et al., 2014, Cowie et al., 2016). The magnitude of this effect is additionally sensitive to the degree of functional coupling as shown by a smaller stopping-interference effect when the prepared response is decoupled (Wadsley et al., 2019; Wadsley et al., 2022b, Jana et al., 2018).

Global inhibitory effects of stopping are typically studied using TMS-derived measures of CME at discrete time points within the estimated time of stopping, however, this methodology has several limitations. First, discrete time points must be determined *a priori* (e.g., Greenhouse et al., 2012), providing poor temporal resolution of response inhibition as it manifests at the effector. Moreover, only one time point can be sampled on a given trial, and MEPs have high trial-to-trial variability, requiring multiple measurements across trials and resulting in long experiments. Additionally, TMS eligibility criteria exclude certain populations of interest and make generalizing the effects of the stopping process more difficult (Rossi et al., 2021). In contrast, surface electromyography (EMG) is safe to use in most populations and offers high temporal resolution, which has revealed informative physiological markers of stopping in task-relevant muscles including within-trial estimates of response inhibition onset (Jana et al., 2020; Raud et al., 2022). The global nature of response inhibition may be detectable in task-irrelevant muscles by measuring tonic muscle activity, since patterns in tonic EMG may be sensitive to global task-dependent changes in corticomotor excitability. Therefore, tonic EMG activity of task-irrelevant muscles may offer a more universally applicable approach to measuring the time course, magnitude, and spatial spread of global inhibition during stopping.

In this study, participants completed a bimanual selective stopping ARI task with their index fingers as responding effectors while maintaining tonic contractions with their pinky fingers to test for signatures of global inhibition in tonic EMG. The EMG activity in the task-irrelevant pinky muscle was compared across go trials, nonselective stop-all trials, and selective stop trials. We hypothesized tonic EMG amplitude in task-irrelevant effectors would transiently decrease during successful and failed stopping compared to go trials, owing to the recruitment of a global inhibitory mechanism. Such a pattern of results could provide valuable information about the time course and extent of the global inhibitory process at the level of the muscles with greater sensitivity than commonly derived behavioral stopping and TMS measurements.

## Methods

### Participants

Twenty participants provided informed consent in accordance with the University of Oregon institutional review board. Potential participants with a history of specific movement disorders were excluded from the study. Out of twenty initial participants, one participant was excluded for poor task performance (less than 25% successful stopping accuracy), and another was excluded for the inability to maintain tonic contractions, which left 18 datasets (9 female, age = 21.6 ± 1.1, all self-reported right handed) for analysis.

### ARI Task

We investigated the effects of the stopping process using a bimanual ARI task, coded in MATLAB 2019a (Figure 1). Participants sat approximately one meter from a computer screen (refresh rate 60 Hz) with their arms supported on armrests, hands extended, and palms facing toward the sagittal midline. The third, fourth, and fifth digits of each hand rested on the surface of the table in front of them in the space between the table and a raised wooden platform of adjustable height. Response buttons were positioned on the elevated platform beneath the medial surface of the left and right index fingers. The response buttons were button switches wired to a MakeyMakey® (Joylabs) connected to the task computer. Participants pressed their pinky fingers downward against the table through abduction, away from the palm, for the complete duration of all task trials.

**Figure 1.**
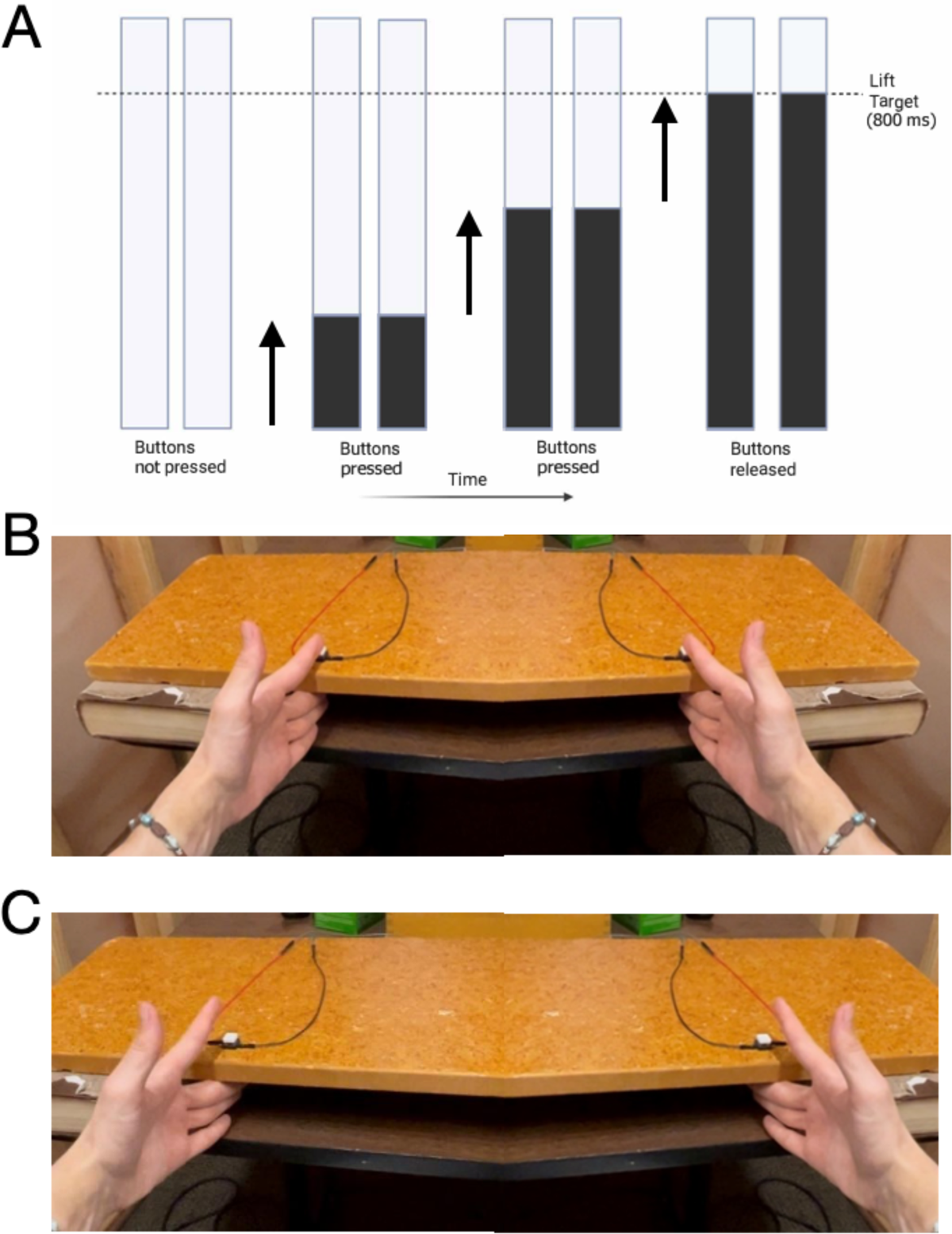
*A.* Stimuli displayed on a computer monitor consisted of two vertical bars that filled from the bottom to the top over the course of a 1 s interval. Participants were instructed to time their responses with when the filling bars intersected a response target line. *B.* Participants tonically contracted the abductor digiti minimi by pressing their pinky fingers downward onto the table surface while pressing both response buttons with their index fingers. *C.* Participants made responses by lifting their index fingers off of the response buttons while maintaining the tonic contraction in the pinky fingers.

The default task display began with two white parallel vertical bars on a gray background. Each task trial started when both buttons on the elevated platform were pressed simultaneously with the index fingers which triggered the bars to begin filling from bottom to top. The bars took one second to fill completely. Participants were instructed to time the lifting of their fingers off of the buttons to the intersection of the filling bars with the target line (800 ms from trial onset, **Figure 1**). Two-thirds of trials were Go trials, during which the left and right bars filled for as long as their corresponding button was pressed, and stopped filling only when the corresponding button was released, i.e. the index fingers were lifted off the buttons.

The remaining one-third of trials were an equal mixture of stop-both, stop-left, and stop-right trials. During stop-both trials, both bars stopped filling automatically after a predetermined stop signal delay (SSD). Participants were instructed to cancel both of their planned index finger lifts while maintaining the contraction with their pinky fingers when the bars stopped. The SSD in this experiment refers to the difference between the target response time and the presentation of the stop signal, and as such is reported as a negative value, where more negative values indicate more time available for stopping. During selective stop-left and stop-right trials, one bar stopped filling at a separate predetermined SSD, while the other bar continued filling to the target. On selective stop trials, participants were instructed to cancel the lift of the finger corresponding to the stopped bar while attempting to lift the remaining finger when its respective bar reached the target line.

SSDs started at −200 ms for stop-both trials and −250 ms for selective stop-left and stop-right trials. Initial SSD values were based on the expected time required for nonselective and selective stopping in healthy young adults (MacDonald et al., 2012). SSDs were adjusted individually for each stop trial type using a staircase procedure (Figure 2). The use of independent SSD staircases was chosen to attain a 50% probability of successful stopping for each stop trial type. The SSD of a particular stop trial type decreased by 50 ms after a failed stop trial (when participants lifted their fingers despite the bar stopping) and increased by 50 ms after a successful stop trial, corresponding to more and less time to stop following failed and successful stop trials, respectively. When participants correctly inhibited a response during a stop trial or lifted the correct buttons within 50 ms of the target line during a go or selective stop trial, feedback was given in the form of the response target line turning green. If the participant responded between 50 and 100 ms of the target line, the response target line turned yellow.

**Figure 2.**
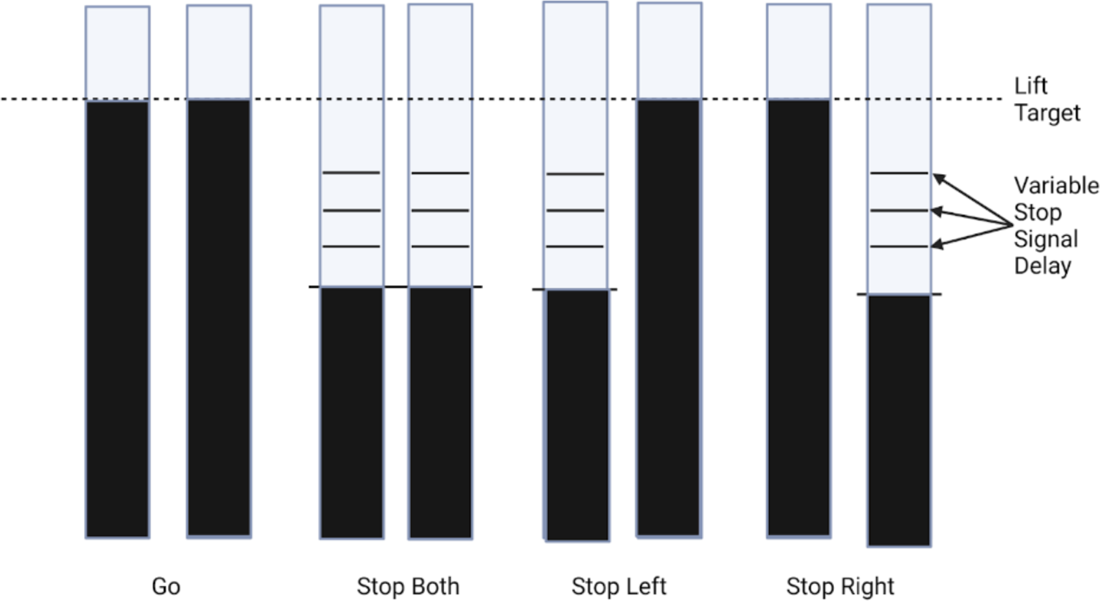
Example task stimuli are shown with each pair of bars corresponding to one of the four types of trials: go, stop-both, selective stop-left, and selective stop-right, respectively. Participants were instructed to lift both index fingers when the bars crossed the target line. If a bar stopped before the response target, participants attempted to cancel the finger lift(s) on the side corresponding to the stopped bar(s).

Finally, if a response was recorded during a stop trial or outside the response window (< or > 100 ms of the response target), the target line turned red. Participants completed 9 blocks of 32 trials (192 Go, 32 stop-both, 32 stop-left, 32 stop-right; 288 trials total), and the order of trial types was randomized for each participant. After each block, participants received feedback about their performance in that block, including their average response time relative to the target line. Participants were encouraged to take breaks between task blocks and instructed to press both buttons when ready to continue.

### Behavioral Metrics

Response time was calculated for all trials by determining the difference between the index finger response time and the target response time (0.8 s). Negative values reflect responses prior to the target and positive values reflect responses after the target. Lift accuracy was calculated for Go and selective stopping trials as the proportion of trials in which the participants responded within 100 milliseconds of the target response time. Stop trial accuracy was calculated as the proportion of trials where the response was successfully withheld. Stop signal reaction time (SSRT), an estimate of the latency of the stopping process was calculated using the integration method with the replacement of go omissions (Verbruggen et al., 2019) separately for the three different types of stop trials. The stopping-interference effect was calculated for the left hand by subtracting the mean response time of the left hand during go trials from the mean response time of the left hand during stop-right trials. The same procedure was employed to calculate the interference effect in the right hand during stop-left trials.

### Electromyography

We recorded surface EMG using bipolar electrodes adhered to the skin above the first dorsal interosseus muscle (FDI) and the adductor digiti minimi (ADM) of both hands. A ground electrode was attached above the styloid process of the left ulna. EMG was sampled at 5,000 Hz, amplified by a factor of 1000, and bandpass filtered (50– 450 Hz; Delsys).

Visualization and analysis of EMG data were performed using the VETA toolbox for MATLAB (Jackson & Greenhouse, 2019). Prior to beginning data collection, participants performed a maximum voluntary contraction (MVC) of each ADM to assess their maximum contractile output. This was done by computing the average maximum EMG amplitude measured during four consecutive 1 s contractions with both pinky fingers. Each participant was instructed to maintain a tonic contraction at 10% of their MVC during the task. Experimenters observed online EMG data traces on a separate monitor positioned adjacent to the stimulus display in order to monitor participants’ contraction and data quality. EMG data was recorded for the duration of each individual trial as well as for 1 second after each trial concluded. Participants completed a set of practice trials (40 trials, 10 of each type) with EMG recording prior to starting the experiment to gain familiarity with the task and to assess EMG data quality, and were allowed to repeat the practice block until they felt confident in completing the task correctly.

For each participant, ADM EMG data were rectified and averaged across trials for each trial type generating 1.8 s duration mean EMG traces for go, stop-both, stop-left, and stop-right trials. These trial average values were then z-scored to determine their variation from baseline during periods of interest and account for between-participant differences in the raw EMG amplitudes. Baseline EMG amplitude was calculated as the average tonic EMG amplitude of the 500 ms preceding the stop signal. The z-scored data of each participant was then rectified across a moving window of 12 sample points, equivalent to a 2.4 ms window (Fisher et al., 2023). Comparisons of EMG across go trials and successful and failed stop trials of all types were performed separately for the left and right ADM. This was done by locking individual trial EMG to stop-signal onset during stop trials and the average SSD on go trials (Jana et al., 2020). Specifically, a 600 ms epoch of interest was defined that started 100 ms before each individual trial’s stop signal for stop trials, and 100 ms before the average SSD for go trials. The duration of this epoch ensured the entire period of the estimated SSRTs was included.

Mean FDI EMG onsets and peak amplitudes were calculated relative to the stop signal for each hand during go trials, failed stop, and successful stop trials of all types and reported in Tables 2 and 3. Onset times were determined using the VETA toolbox ‘findEMG.m’ function (Jackson & Greenhouse, 2019). In brief, the FDI EMG onset times were determined to be the first time points at which the rectified EMG signal exceeded two standard deviations of the mean within a trial.

### Statistical Analyses

We tested for differences in SSRT and stop accuracy across stop trial types using a 1 × 3 (stop-both, stop-left, stop-right) repeated measures (RM) ANOVA with Bonferroni corrected post-hoc paired *t*-tests. We tested for differences in response times and response accuracies using 2 × 3 RM ANOVA with the factors Hand (Left, Right) and Trial Type (Go, Failed Stop-Both, Selective Stop Response). The magnitude of the stopping-interference effects during selective stop trials was compared between hands using a paired sample *t*-test.

To test for suppression of the tonic EMG signal during the stopping epoch, time series analyses were conducted using pairwise t-tests at each of 3000 time points of interest across trial types. These time points correspond to the previously mentioned 600 ms epoch. A false discovery rate (FDR) correction with an adjusted alpha level of 5% was used to account for multiple comparisons in each case (Benjamani and Hochberg, 1995). Specific comparisons of interest included go vs successful stop-both, go vs successful stop-left, and go vs successful stop-right trials. Additional analyses compared failed stopping between failed stop-both, failed stop-left, and failed stop-right trials using the same approach. Data from the left and right ADM were analyzed separately in all cases.

## Data Availability

All data and analysis code are available for download via Open Science Framework (osf.io/gkeb3).

## Results

### Behavioral Data

Behavioral metrics are presented in Table 1 as mean ± standard deviation. High go accuracy in conjunction with stopping accuracies close to 50% across all stop trial types indicated that participants performed the task correctly and did not adjust their strategies to anticipate stop signals. There was a significant main effect of trial type for SSRT [F(2,17) = 3.39, *P =* 0.0319]. Post-hoc t-tests showed stop-right SSRT values were significantly longer than stop-both SSRTs (*t* = 2.62, *P =* 0.0927), and stop-left SSRTs were longer than stop-both SSRTs (*t* = 2.43, *P =* 0.0137). The difference between stop-left and stop-right SSRTs did not reach significance (*t* = 0.34, *P =* 0.371). There was no effect of trial type on stopping accuracy for the stop-both, stop-right, and stop-left conditions [F(2,17) = 0.52, *P =* 0.603].

**Table 1.** Behavioral metrics of interest Mean ± std.

| Trial Type | Go Left | Go Right | Stop-Both | Stop-Left | Stop-Right |
| --- | --- | --- | --- | --- | --- |
| Response time (ms) | 16 ± 5 | 12 ± 5 | | 111 ± 29 | 106 ± 32 |
| Interference effect (ms) | | | | 95 ± 27 | 94 ± 33 |
| Failed Stop RT (ms) | | | -11 ± 8 | -15 ± 26 | -3 ± 28 |
| SSRTs (ms) | | | 287 ± 52 | 310 ± 46 | 314 ± 47 |
| SSD (ms) | | | -237 ± 45 | -258 ± 38 | -262 ± 55 |
| Stop Accuracy (%) | | | 48 ± 2 | 50 ± 2 | 49 ± 3 |
| Lift Accuracy (%)* | 88 ± 6 | 91 ± 5 | | 49 ± 16 | 51 ± 21 |
| Failed Stop Trial Lift Accuracy (%)* | | | 98 ± 2 | 86 ± 11 | 84 ± 17 |

Abbreviations: RT, response time; SSRT, stop signal reaction time; SSD, stop signal delay. \* indicates proportion of responses ± 100 ms of lift target.

**Table 2.** FDI EMG event times relative to the stop signal (mean SSD for Go trials) and number of partial responses (mean ± std)

| Trial Type | Go | FSB | FSL | FSR | SSB | SSL | SSR |
| --- | --- | --- | --- | --- | --- | --- | --- |
| Left FDI EMG onset (ms) | 114 ± 14 | 44 ± 26 | 108 ± 33 | 148 ± 32 | 110 ± 31* | 166 ± 34* | 263 ± 36 |
| Left FDI EMG peak (ms) | 188 ± 15 | 142 ± 24 | 213 ± 55 | 253 ± 47 | 142 ± 34*<sup>c</sup> | 205 ± 73*<sup>c</sup> | 352 ± 56 |
| Left FDI partial responses (#) | | | | | 5.1 ± 3.6 | 6.4 ± 2.9 | |
| Right FDI EMG onset (ms) | 111 ± 17 | 49 ± 25 | 103 ± 21 | 143 ± 25 | 107 ± 37* | 224 ± 41 | 203 ± 55* |
| Right FDI EMG peak (ms) | 178 ± 20 | 129 ± 32 | 174 ± 42 | 252 ± 53 | 138 ± 35*<sup>c</sup> | 300 ± 46 | 380 ± 87*<sup>c</sup> |
| Right FDI partial responses | | | | | 6.3 ± 3.4 | | 6.9 ± 4.4 |

Abbreviations: FSB: Failed stop-both; FSL: Failed stop-left; FSR: Failed stop-right; SSB: Successful stop-both; SSL: Successful stop-left; SSR: Successful stop-right); \* indicates partial response trials; <sup>c</sup> cancel time

A 2 × 3 RM ANOVA with the factors Hand (Left, Right) and Trial Type (Go, Failed Stop-Both, Selective Stop Response) showed no difference in response time for the main effect of Hand [F(1,17) = 0.13, *P =* 0.724], and a main effect of trial type [F(2,17) = 164.7, *P <* 0.001], with no interaction between factors []. Post-hoc analyses revealed that stop-right response times (left index responses) were significantly longer than go trial left hand response times (*t* = 14.3, *P <* 0.001), and failed stop-both left hand response times (*t* = 12.7, *P <* 0.001). Stop-left response times (right index responses) were significantly longer than go trial right hand response times (*t* = 11.4, *P <* 0.001), and failed stop-both right hand response times (*t* = 8.8, *P <* 0.001). Right hand failed stop-both responses were shorter than go responses (*t* = 4.6, *P <* 0.001), but the difference between left hand failed stop-both responses and go responses did not reach significance after correcting for multiple comparisons (*t* = 2.1, *P =* 0.0239, uncorrected). There was no difference in the magnitude of the stopping-interference effect between hands (*t* = 1.96, *P =* 0.0668).

The majority of lifts occurred within ±100 ms of the target, corresponding to high response accuracies (≥84%) across go trials and failed stop trials. Lift accuracies were lower among responding hands during successful selective stop trials (≥48%), but this was an expected manifestation of the interference effect. A 2 × 3 RM ANOVA with the factors Hand (Left, Right) and Trial Type (Go, Failed Stop-Both, Selective Stop Response) revealed a significant effect of trial type on lift accuracies [F(2,17) = 167.9, *P <* 0.001], with no difference between hands [F(2,17) = 0.131, *P =* 0.718], and no interaction between factors [F(2,17) = 0.087, *P =* 0.917].

Post-hoc t-tests showed that right hand selective stop response lift accuracy was lower than go (*t* = 9.1, *P <* 0.001), and failed stop-both lift accuracy (*t* = 9.7, *P <* 0.001). Left hand selective stop response lift accuracy was lower than go (*t* = 8.4, *P <* 0.001) and failed stop-both lift accuracy (*t* = 9.1, *P <* 0.001). There was no difference in lift accuracy between failed stop-both and go trials in the left (*t* = 0.1, *P =* 0.466) or the right hand (*t* = 0.2, *P =* 0.413).

### EMG Data

FDI EMG onsets and peak amplitudes were consistent with the behavioral measures and are presented in Tables 2. Of note, FDI EMG onset and peak times were similar to those found in the ADM EMG data when comparing go and failed stop trials, although the nature of the task design led to relatively few partial responses being recorded for each stop trial type.

Tonic ADM EMG amplitudes during failed stop-both, failed stop-left, and failed stop-right trials exhibited an interval of divergence from go trials characterized by a clear transient dip. To determine the onset and offset of this interval for each type of failed stop trial in each hand, we conducted a post-hoc analysis. We identified the longest interval in which more than 90% of timepoints showed a significant (FDR corrected) difference between failed stop and go trials. During stop-both trials, the left ADM exhibited significant decreases in tonic EMG output as compared to go trials from 149 ms to 194 ms following the stop signal, while the right ADM exhibited significant decreases in tonic EMG output from 150 ms to 207 ms (Figure 3). During Stop-Left trials, the left ADM exhibited significant decreases in tonic EMG output from 152 ms to 205 ms, while the right ADM exhibited significant decreases in tonic EMG output from 150 ms to 193 ms (Figure 4). During Stop-Right trials, the left ADM exhibited significant decreases in tonic EMG output from 155 ms to 210 ms, while the right ADM exhibited significant decreases in tonic EMG output from 145 ms to 209 ms (Figure 5). Overall, these results describe an average period of suppression during failed stopping as compared to go trials from 150 ms to 203 ms.

**Figure 3.**
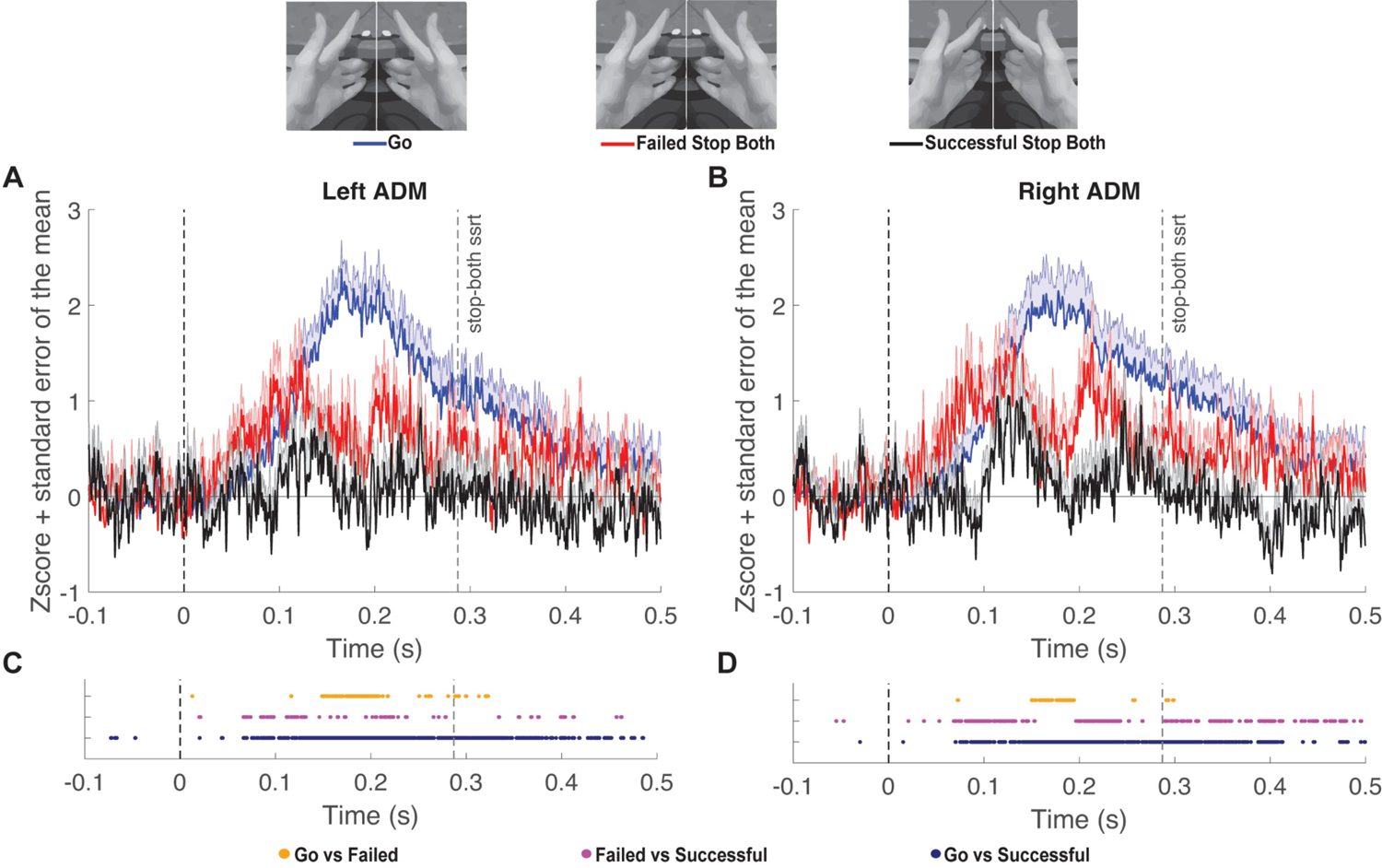
*A*. Z-scored tonic left ADM EMG amplitude (mean + sem) comparing Go trials locked to the mean SSD and Stop-Both trials locked to the stop signal at time point zero. Group mean SSRT is denoted by the rightmost vertical dashed line. *B*. Same as A for the right ADM. *C*. Significant FDR corrected comparisons (*P <* 0.05) for all time points of interest for the left ADM. *D*. Same as C for the right ADM.

**Figure 4.**
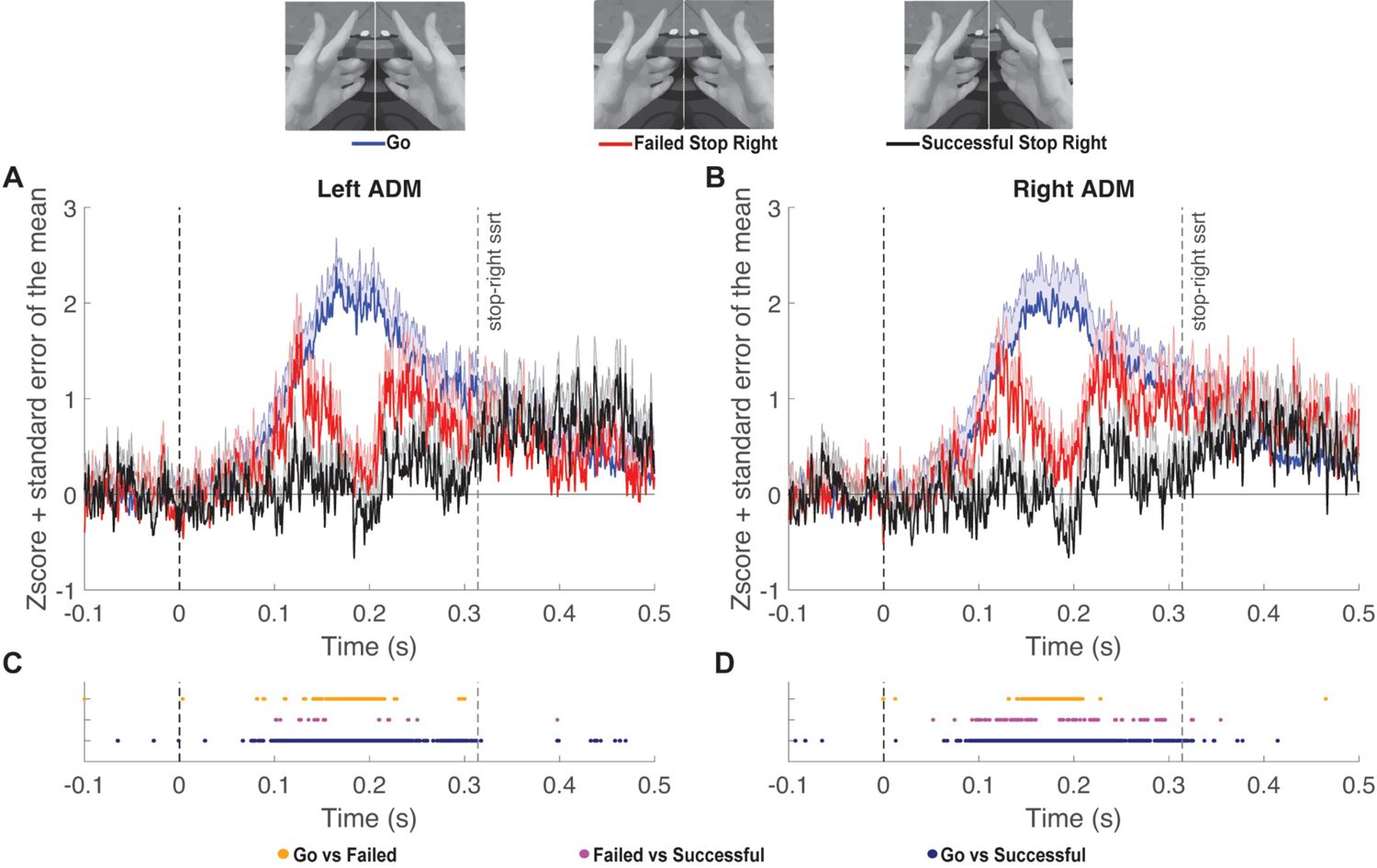
*A*. Z-scored tonic left ADM EMG amplitude (mean + sem) comparing Go trials locked to the mean SSD and Stop-Right trials locked to the stop signal at time point zero. Group mean SSRT is denoted by the rightmost vertical dashed line. *B*. Same as A for the right ADM. *C*. Significant FDR corrected comparisons (*P <* 0.05) for all time points of interest for the left ADM. *D*. Same as C for the right ADM.

**Figure 5.**
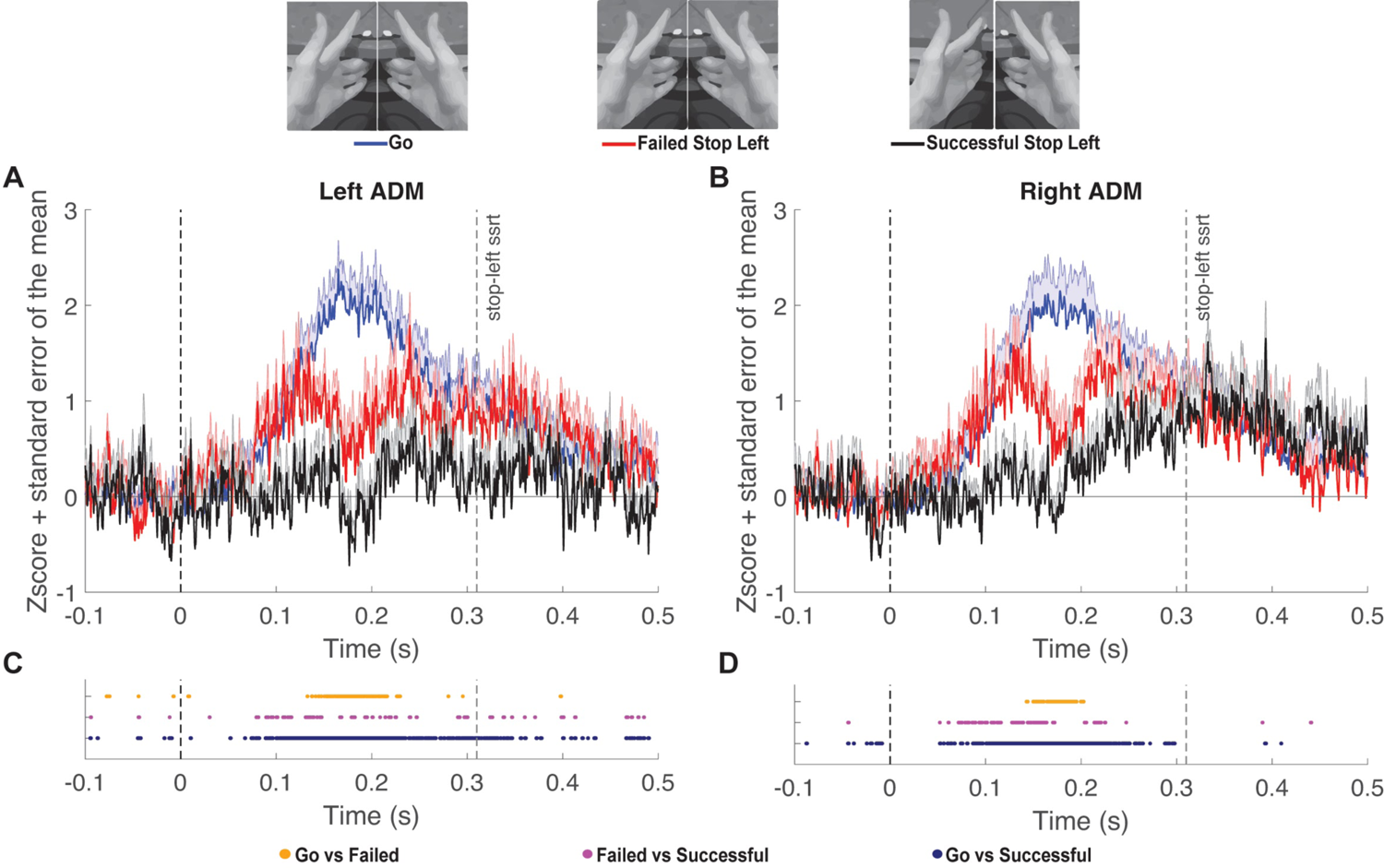
*A*. Z-scored tonic left ADM EMG amplitude (mean + sem) comparing Go trials locked to the mean SSD and Stop-Left trials locked to the stop signal at time point zero. Group mean SSRT is denoted by the rightmost vertical dashed line. *B*. Same as A for the right ADM. *C*. Significant FDR corrected comparisons (*P <* 0.05) for all timepoints of interest for the left ADM. *D*. Same as C for the right ADM.

Our design permitted us to compare between three types of failed stop trials: stop both, selective stop left, and selective stop right. The contrast across these conditions is unique because the behavioral outcomes are matched, i.e. both index fingers are lifted in all three cases, and only the context differs. Differences in the pattern of tonic EMG between the bimanual and selective unimanual failed stop trials could suggest the operation of different inhibitory mechanisms. However, after FDR correction, there were no time points identified in which tonic ADM EMG amplitudes differed across the three types of failed stop trials (Figure 6).

**Figure 6.**
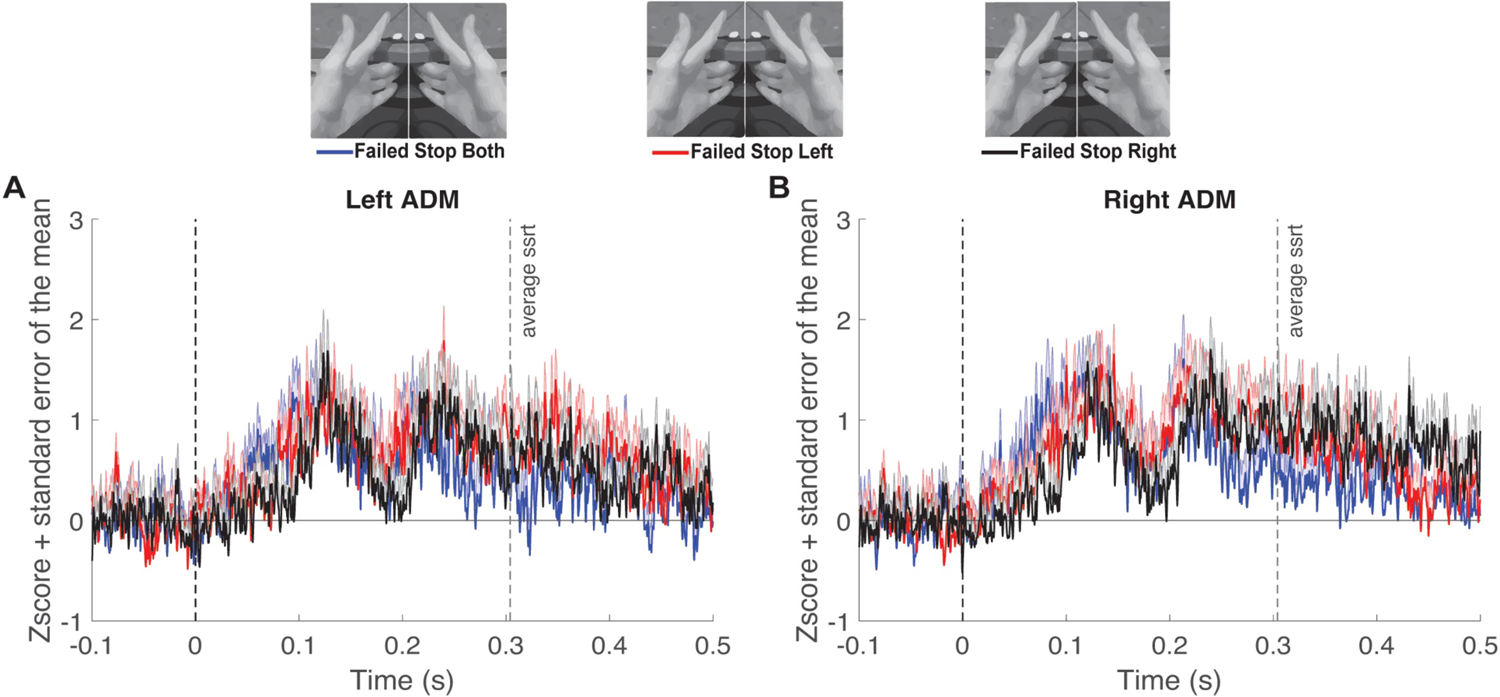
*A*. Z-scored tonic left ADM EMG amplitude (mean + sem) comparing Failed Stop trials of all types locked to the stop signal at time point zero. Mean SSRT is denoted by the rightmost vertical dashed line. *B*. Same as A for the right ADM. FDR corrected comparisons (*P <* 0.05) were performed, but showed no significant differences between trial types at any time point.

## Discussion

We analyzed the EMG of tonically contracting, nonresponding muscles during a selective ARI task to determine whether EMG can provide a continuous and temporally sensitive measure of global motor inhibition. We discovered a transient period of tonic EMG suppression during failed stopping compared to going between 150 and 203 ms following the stop signal. The timing of this pattern is consistent with the hypothesized downstream effects of the hyperdirect pathway for interrupting ongoing motor responses. Moreover, this pattern was highly consistent in both hands regardless of whether the stop signal was bimanual or selective (unimanual). This further supports the existence of a nonselective, potentially global, stopping mechanism that suppresses muscle activity for approximately 50 ms.

Our approach to measuring tonic EMG in the context of selective stopping is novel. However, previous studies examined EMG characteristics of the stopping process in the absence of tonic EMG activity. Raud and Huster (2017) observed partial EMG bursts (referred to as “subthreshold EMG” in their paper) in the responding effector during successful stopping, indicating the EMG activity associated with the initiation of a response can be rapidly terminated. They interpreted the peak of these partial EMG bursts as a physiological marker of when the stop process reached the muscle, approximately 147 ms after the stop signal during a reactive inhibition task and at 152 ms during a proactive inhibition task. Other subsequent studies used the partial EMG response peak to estimate the latency of the onset of inhibition, or ‘cancel time,’ during nonselective (Jana et al., 2020; Raud et al., 2020a, 2020b, & 2022) and selective stopping (Wadsley et al., 2022b; Salomoni et al., 2024). The observed onset of inhibition in task-relevant muscles in these previous studies closely matches the onset of EMG suppression we observed during failed stopping in the current study.

Using the tonic EMG trace, we were also able to estimate the release of inhibition based on the time at which EMG activity on failed stop trials returned to a level similar to that of go trials. The offset of this inhibition period has been more difficult to estimate in past studies. Studies incorporating TMS to measure CME modulation during stopping often only focused on one or a handful of time points in which they reported evidence of global inhibition. Many of these studies reported measurable decreases in MEP amplitudes within the same time frame following the stop signal in which we observed EMG suppression (Badry et al., 2009; Macdonald et al., 2014; Cowie et al., 2016). While individual studies reported measurable decreases in CME as early as 140 ms (Coxon et al., 2006) and as late as 220 ms (Majid et al., 2012) following the stop signal, the differences between these values and our reported values are small and may reflect differences in the behavioral tasks or the influence of inhibition operating at the cortical vs muscular level.

Interestingly, nearly all past studies that reported transiently reduced CME during successful stop trials report no reduction during time-matched stimulation on failed stop trials (Badry et al., 2009; Majid et al., 2012, Greenhouse et al., 2012), or non-significant reductions (Cai et al., 2012). This suggests tonic EMG is more sensitive than TMS to inhibition in the context of ongoing responses, at least in the case of failed stopping. Alternatively, the observed inhibition during failed stopping may be a product of our task design and the proximity of the tonically contracted ADM to the responding FDI. While we identified many timepoints with significantly lower ADM activation during successful stopping compared to go trials and failed stop trials, the patterns were less consistent and were not subjected to the same interval testing. One pattern of note is that during the 150 ms to 203 ms period of suppression in failed stop trials, there was a considerable reduction in the density of timepoints showing significant differences in EMG amplitude between successful and failed stopping. This pattern is most noticeable in Figures 3B, 4B, and 5A & B, and indicates tonic EMG amplitudes during failed stop trials were reduced to values in a similar range to those of successful stop trials within this interval of transitory inhibition before rebounding to activation levels more similar to go trials. This pattern could suggest that the motor system is inhibited to a similar extent for both types of stop trials, and the influence of inhibition on background activity is most visible within the context of a response.

Notably, we were unsuccessful in previous attempts to measure markers of inhibition in tonic EMG using two versions of unimanual reactive stop signal tasks. Specifically, we did not observe fluctuations in tonic EMG in a non-responding hand during the performance of simple or choice versions of a standard stop task (Fisher et al., 2023). We speculate this may have been due to the decoupling of the hands during those tasks. By contrast, the bimanual and symmetrical nature of the ARI task employed here encourages participants to couple their hands to respond, which may explain why the patterns of inhibition described above were detectable in the tonic EMG (McDonald et al., 2021; Wadsley et al., 2022).

Global motor inhibition is hypothesized to be governed by the hyperdirect pathway, a fast cortico-basal ganglia pathway that bypasses the striatum via the subthalamic nucleus (STN) (Aron & Poldrack, 2006), and is strongly implicated in reactive stopping (Chen et al., 2020). Historically, behavioral stopping has been characterized as a horse race between a Go and Stop process, with the faster process determining the behavioral outcome (Logan and Cowan, 1984), but more recent analyses of stopping suggest stopping may be a two-part process in which a nonselective pause temporarily globally inhibits the motor system, followed by a selective cancel phase which revises or completely cancels active motor programs. In rodent models (Schmidt and Burke, 2017) and in humans (Diesberg and Wessel, 2021) the pause phase is theorized to be initiated by the hyperdirect pathway, while the cancel phase reflects the action of the slower indirect pathway. In this description of stopping, the pause phase is triggered following all “salient” stimuli including the presentation of the stop signal, resulting in a transient episode of inhibition during successful as well as failed stopping in task paradigms such as the SST and ARI task. Corroborative physiological evidence suggests a global pause mechanism is engaged during the early parts of both selective and non-selective stopping (Raud et al., 2020; Wadsley et al., 2023).

There are hints of a potential cancel process in our data. Although we did not observe sustained levels of significance comparable to the 150 ms to 203 ms interval, we visually identified a second decrease in tonic EMG amplitude around 250-300 ms after the stop signal during successful and failed stop trials relative to go trials (Figures 3-6 for successful stopping). We speculate that such a second wave of inhibition is consistent with recently proposed descriptions of stopping as a two part pause-then-cancel process (Schmidt and Burke, 2017; Diesberg and Wessel, 2021). In particular, the time period of this second wave coincides with the purported role of electroncephalography-dervied P300 signature in response cancellation (Wessel & Aron, 2015). Future studies can elucidate the potential separable influence of pause and cancel processes on tonic muscle activity by using a stimulus-selective response inhibition paradigm (e.g., Tatz et al., 2021; Wadsley et al., 2023).

The identified period of inhibition across stop trial types combined with the temporal alignment with previously characterized physiological markers of stopping suggests our observed decrease in tonic EMG reflects the hyperdirect pathway’s influence on both task-relevant and irrelevant muscles during failed stopping. Our approach offers several advantages over others that have been used to measure such markers of motor inhibition. EMG is noninvasive and is a less exclusionary option for assessing physiological mechanisms involved in stopping compared with TMS or MRI. EMG also possesses excellent temporal resolution compared with those techniques, and individual trial measurements of tonic EMG activity may provide further insights into trial-to-trial variability. Therefore, this approach may be particularly appropriate for assessing hyperdirect pathway function in patients who suffer from diseases associated with basal ganglia dysfunction such as dystonia, chorea, ballism, or Parkinson’s disease.

### Limitations

Sustaining a tonic contraction in the ADM muscle may have influenced the manner in which participants performed the ARI task. However, our data are very similar to previous studies that have used the task without additional ongoing muscle contractions (Wadsley et al., 2023).

Moreover, the staircase procedure for setting the SSD would adjust for factors that could potentially interfere with the stopping process. Inhibition in the task-irrelevant ADM was not as apparent during successful as during failed stop trials. This was likely because successful stop trials include a mixture of trials in which there are partial EMG responses and trials in which the EMG never initiates. The tonic ADM EMG exhibited an apparent increase coupled to FDI response EMG onset during Go and Failed Stop trials. This coupled increase in EMG was not necessarily expected but may have exaggerated the differences between going and failed stopping in this particular version of the task. The slight elevation in the tonic ADM EMG activity coupled with the responding FDI EMG may have been sufficient to reveal inhibition that remained hidden during successful stop trials. Furthermore, the EMG burst associated with response execution on failed stop trials may provide sufficient homogeneity for inhibition to be detectable, given the low likelihood of ‘trigger failures’ and relatively narrow distribution of onset times. Relatedly, failed stopping across all trial types (stop-both, stop-left, and stop-right) involved bimanual responses resulting in similar EMG patterns across these trial types. We only measured intrinsic hand muscles, and future studies may be able to examine this effect in coupled muscles further apart from the primary responding effector to evaluate the spatial spread of inhibition in tonic EMG. Future studies will also be able to evaluate whether our results generalize to other versions of stop tasks such as a unimanual ARI task.

## Conclusion

In this study, we observed an electromyographic marker of inhibition acting at the periphery in a non-responding muscle during action stopping. This measure can provide unique insight into the onset timing, magnitude, spatial spread, and duration of the underlying neural mechanisms. Our data are consistent with the engagement of a global ‘pause’ mechanism that inhibits muscle activity starting approximately 150 ms after the stop signal and persisting for approximately 50 ms. This finding has immediate relevance to other studies of response inhibition as well as other behaviors associated with widespread transient motor system inhibition (e.g. detection of unexpected events). Our findings may be useful in contextualizing abnormalities in the dynamics of inhibitory processes acting on the motor system, and the non-invasive nature of our approach lends itself well to applications in a wide variety of clinical and healthy populations.

